# Ability of a Structural World Model to Detect Cryptic Pockets from Apo Structure

**DOI:** 10.64898/2026.09.21.752781

**Authors:** Ryan Shihabi, Brent Vaughan, Sharief Taraman

## Abstract

Cryptic pockets are druggable sites that are absent in a protein’s resting structure and form only upon backbone rearrangement, posing a challenge for detection from static apo structures. Existing methods rely on generating open conformations through sampling or complex prediction, limiting applicability. Here we present a novel detector that reads cryptic pockets directly from a structural world-model latent representation of a single apo structure, without conformational sampling or external pocket finders.

Evaluated on CryptoBench and CryptoBank datasets, the method localizes cryptic sites with top-1 accuracy as high as 0.848 and top-5 accuracy exceeding 0.93, and successfully recovers an allosteric site on held-out WRN helicase structures. The detector complements existing approaches and improves with training data scale. These findings suggest that structural world-model latents encode conformational flexibility, enabling effective cryptic pocket identification from apo structures alone, facilitating drug discovery on challenging targets.

Benchmarked against the co-folding engine OpenDDE, the detector finds pockets directly rather than by first predicting a bound complex: on 190 targets held out of both training sets it recovers 119 sites OpenDDE misses, against 2 in the other direction, and at top-5 co-folding’s recovered sites are a subset of ours. It runs on any structure from the apo coordinates alone, including the mmCIF-only entries all recent depositions carry, and its accuracy keeps climbing as the training corpus grows. The intended use is prospective cryptic-site nomination on the targets that sequence and static structure leave without a starting point.

## 1. Introduction

### 1.1 The Site That Is Not in the Structure

Most drugs act at a pocket, and most pocket finders identify such pockets directly from the resting structure. However, a cryptic pocket challenges this approach, as it is closed, occluded, or simply absent in the apo conformation and only forms after the backbone rearranges, often upon ligand binding. Detecting such a site is therefore not a matter of refining surface geometry, but instead requires identifying which residues will line a site that has not yet opened. Many targets that appear featureless or undruggable in their crystal structures actually harbor these cryptic sites, making their detection central to expanding the set of druggable proteins.

The key evidence that a static method would require, the open conformation, is inherently unavailable, as it is precisely what one aims to discover. For most targets, no companion structure exists to reveal the open state, so any method that relies on the open conformation, a bound ligand, or a solved holo structure cannot be applied to the targets that need it most. Therefore, the central challenge is to identify cryptic residues using only the apo coordinates, for any protein.

### 1.2 Approach

Traditionally, detecting a cryptic site requires generating the open state, either by sampling conformations through molecular dynamics or elastic-network models, or by predicting a bound complex using co-folding models. In contrast, our world model encodes this information internally: trained on structural context, its latent representation has learned where the backbone can move and where a site can open. As a result, the conformational signal that other methods must generate at inference is already present in the representation of the closed apo structure. Thus, detection requires no external feature space; from a single static structure, without ensembles or language models, the latent identifies the site that other methods reconstruct by producing the open state.

In the approach contained herein, cryptic sites are read from a single world-model latent [2, 10], used directly and left unspecified here (see Note). Each residue receives one latent vector, and the same vector that places a residue in structural context also scores whether it lines a site that opens.

By world model we mean the term in its specific sense [2], not a loose synonym for a large foundation model. A world model supports planning: it maps observation to state, predicts how an action changes that state, and scores the reward that follows, so the outcome of an action can be anticipated before it is taken. Its representation is created by native molecular interaction, so the latent encodes how molecules actually organize and come into contact. That is how the latent here should be read: it is not a sequence or structure embedding that happens to separate cryptic residues, but a state from which physical relations are predicted, and the cryptic score is one such relation read back out. The model itself, why it earns the term, and its evaluation as a world model are set out for HI-JEPA [10]; here we report one capability read from that state.

From this latent, a candidate pocket is constructed as a grown, disjoint set of residues, seeded at a high-scoring residue and expanded according to an explicit rule, rather than being constrained to a fixed-radius sphere (Figure 1). This approach allows the predicted pocket to better match the expected shape defined by the evaluation metric. No external tools or predictions are incorporated into the output: no separate pocket finder is used, and no external predictions are appended to our results.

**Figure 1:**
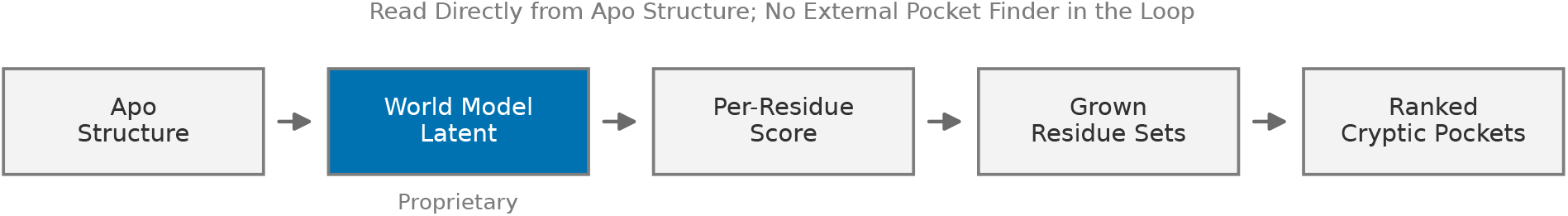
The detector, end to end. An apo structure is read into a world-model latent (proprietary, see Note), scored per residue, grown into disjoint residue sets under an explicit rule, and ranked into candidate cryptic pockets. The representation is read directly; no external pocket finder is in the loop.

### 1.3 Contributions

- **Detection from a World-Model Latent, Read Directly**. The representation is scored per residue and grown into pockets, with the conformational signal carried in the latent rather than sampled at inference, and it generalizes to targets with no solved open-state structure (Sections 5, 6).
- **Ranking the Site First**. Ensemble detectors are able to recover cryptic sites, but they do not reliably rank the true site as the top candidate [11]. In contrast, because our method uses the same per-residue latent score to both identify and rank candidate regions, the true site is often placed at rank 1, rather than simply being present among the top few (Section 8.1).
- **A Held-Out Target**. On WRN helicase, held out of training, the allosteric site is recovered at rank 1 on three independent apo forms (Section 6.1).
- **Direct Detection, Not Ligand-First.** Co-folding reaches a pocket only by predicting its bound complex; our detector finds it directly from apo coordinates, and at top-5 co-folding’s recovered sites are a subset of ours (Section 8.2).
- **Accuracy That Scales with Data**. The detector improves with the size of the training corpus, and that growth is the primary route to a better detector (Section 7).

**Table 1.** Summary of Principal Results. The first five rows report absolute performance; the last three are head-to-head comparisons, each scored under the baseline’s own rule and annotated with the rank compared. Paired values are top-1 / top-5, the WRN row gives the pocket rank on each of three apo forms, and a dash marks a row with no baseline comparison.

| Result | Corpus | This Work | Baseline | Section |
| --- | --- | --- | --- | --- |
| Localization, overlap rule | CryptoBench (231) | 0.848 / 0.952 | - | 5.1 |
| Localization, overlap rule | CryptoBank (286) | 0.846 / 0.990 | - | 5.1 |
| Per-residue detection (AUC) | CryptoBench (231) | 0.8465 | - | 5.2 |
| Exact residue set, Jaccard rule | CryptoBench (231) | 0.597 / 0.628 | - | 5.2 |
| Held-out allosteric site | WRN helicase | 1 / 1 / 1 | - | 6.1 |
| Recovery vs Lacuna, default | CryptoBench (231) | 0.697 (top-1) | 0.556 (top-5) | 8.1 |
| Recovery vs Lacuna, PLM | CryptoBench (231) | 0.697 (top-1) | 0.661 (top-5) | 8.1 |
| Unique sites vs OpenDDE | 190 held-out | 119 (top-1) | 2 (top-1) | 8.2 |

## 2. Related Work

Cryptic-pocket detection falls into three main families. Conformational-ensemble methods generate open-like conformations, by molecular dynamics or elastic-network sampling, and then score pockets across the resulting ensemble; fpocket [3] is a widely used geometric detector that scores cavities on a single structure and is often applied conformer by conformer, and Lacuna [7], the open-source baseline we compare against, wraps the full ensemble pipeline into one tool, generating a conformational ensemble from an input structure, detecting pockets in every conformer, clustering the corresponding pockets across the ensemble to surface sites that appear only transiently, and ranking the resulting sites with a learned model. A second family of supervised residue classifiers predicts cryptic residues directly from combinations of sequence and structure features; geometry-based site finders such as P2Rank [4] rank surface pockets on a single structure, while learned single-structure predictors such as PocketMiner [12], a graph network trained to predict where pockets open, and DeepPocket [13], a 3D convolutional network that rescores geometric cavities, score sites from one conformation. More recently, folding and co-folding models [1] are read for pocket opening through their prediction of a bound complex, and proprietary drug-design engines such as IsoDDE [9] extend co-folding to ligandable-pocket identification.

Our detector reads a single learned world-model latent over the apo coordinates, scored per residue and grown into pockets, with the conformational signal carried in the representation rather than sampled at inference. It differs from the ensemble family in not sampling conformations at test time, from the classifier family in reading a learned world-model latent rather than hand-specified features, and from the co-folding family in requiring neither a partner nor a predicted complex. Where we report any of them as a baseline we run it at its own strongest setting rather than quoting a published headline, since the hit rule, the pocket size, and the split each move the number.

## 3. The Detector

### 3.1 Terms

- **Residue Latent**, the single vector the world model assigns to one residue of the apo structure.
- **Candidate**, a grown, disjoint set of residues seeded at a high-scoring residue and expanded under a fixed rule to a chosen size.
- **Extraction Cell**, the seed-count and grow-radius pair that fixes candidate shape, selected on training data under the rule being reported.
- **Rule**, the pocket-level hit criterion: overlap (at least three residues shared with the annotated site), Jaccard (intersection over union at least 0.25), or DCA (predicted center within four angstroms of a site atom). The three measure different things and are never blended.

### 3.2 Construction

The detector processes a single structural world-model latent over the apo chain, assigning a latent vector to each residue; the inputs and internal structure of the model are proprietary (see Note). Each residue is scored individually, and candidate pockets are seeded at high-scoring residues, then grown according to a fixed rule to a specified size. Geometric non-maximum suppression is applied to prevent reported pockets from overlapping, ensuring that each rank corresponds to a distinct site. To stabilize the ranking, the detector pools scores from a fixed ensemble of independent scoring seeds, rather than from different conformations, which reduces run-to-run variance in the readout without introducing bias; the pooled score is used to rank the candidates. The apo structure is read only once, with no conformational sampling. In the configuration used throughout this work, the extraction cell is selected on training data under the overlap rule and capped a priori at the scale a drug-binding site lines. These extraction settings are held constant across all datasets, so a single configuration generates all reported results.

## 4. Data and Evaluation Protocol

We evaluate two public cryptic-pocket corpora and one held-out case. CryptoBench [5] provides paired apo and holo structures with annotated cryptic residues; we score its held-out test split of 231 chains, with three large multi-chain assemblies that fail structural preprocessing counted as misses rather than dropped, so the denominator is the full 231. From CryptoBank [6] we score a separate 286-chain subset, and the WRN helicase case (Section 6.1) uses public ligand-bound structures as ground truth. Every chain is scored from the apo structure alone, with no holo structure and no bound ligand entering the prediction.

Two points govern how the numbers should be read. First, targets are held out by protein and not by chain: a held-out chain of a protein already seen in training is not a held-out target, so only proteins entirely absent from training count as held out. Second, every reported number states its hit rule and denominator. Overlap only rises as candidates grow larger, so pocket sizes are matched before any cross-method comparison, and DCA is scored the same way as the baselines, the predicted center against the nearest site atom, giving a top-1 of 0.649 (about 0.03 of spread across seeds). Per-residue AUC is estimated by bootstrap across seeds.

## 5. Detection Accuracy

### 5.1 Localization on CryptoBench and CryptoBank

Using only the apo structure and applying the pocket-level overlap rule, the detector localizes the cryptic site with the following performance metrics.

| Corpus | n | Top-1 | Top-3 | Top-5 |
| --- | --- | --- | --- | --- |
| CryptoBench Test | 231 | 0.848 | 0.931 | 0.952 |
| CryptoBank Subset | 286 | 0.846 | 0.933 | 0.990 |

Localization is already strong at rank 1 and has saturated by rank 5 on both corpora (Figures 2 and 3), so the cryptic site is nearly always among the first few pockets the detector returns.

**Figure 2:**
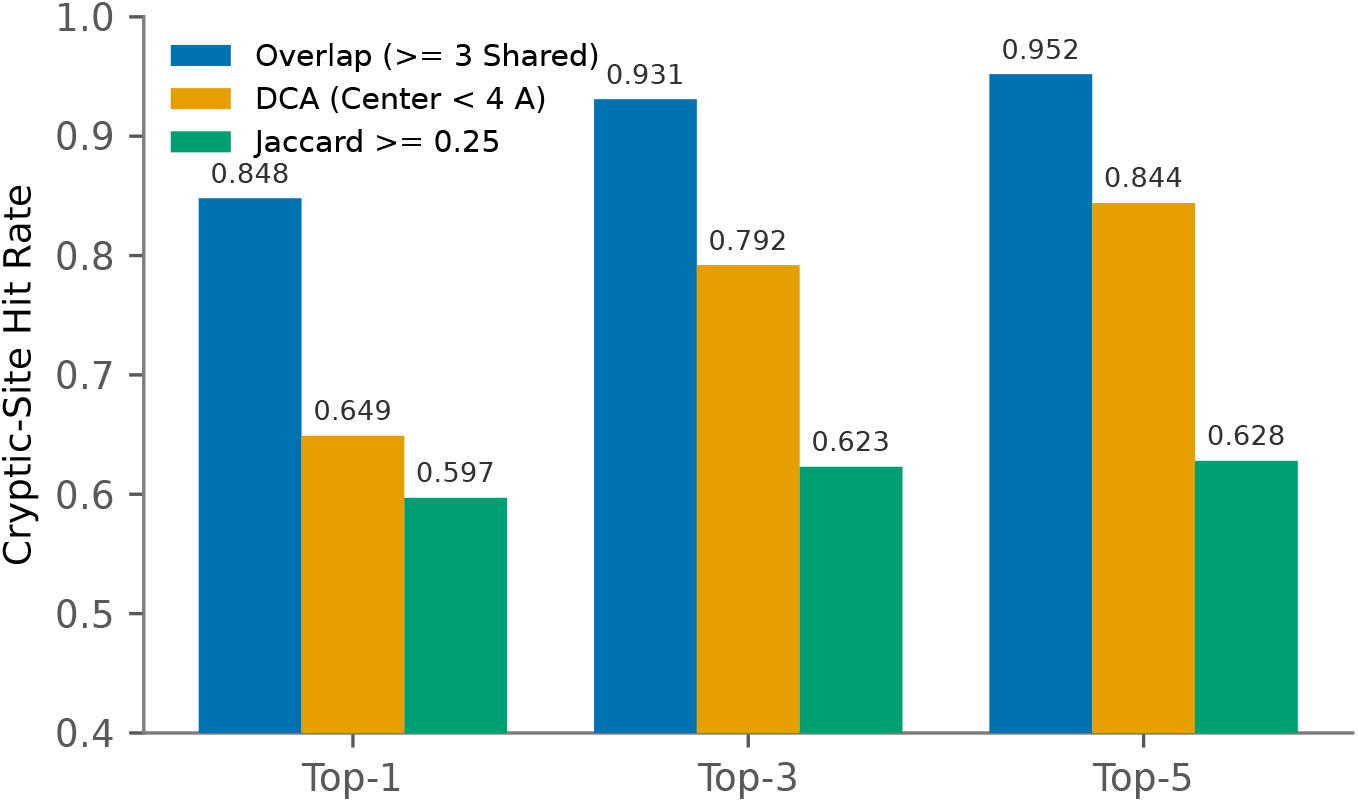
CryptoBench detection across rules. Cryptic-site hit rate on CryptoBench at ranks 1, 3 and 5 under three rules: the pocket-level overlap rule, the DCA center-distance rule (center within four angstroms of a site atom), and the strict Jaccard rule (at least 0.25).

**Figure 3:**
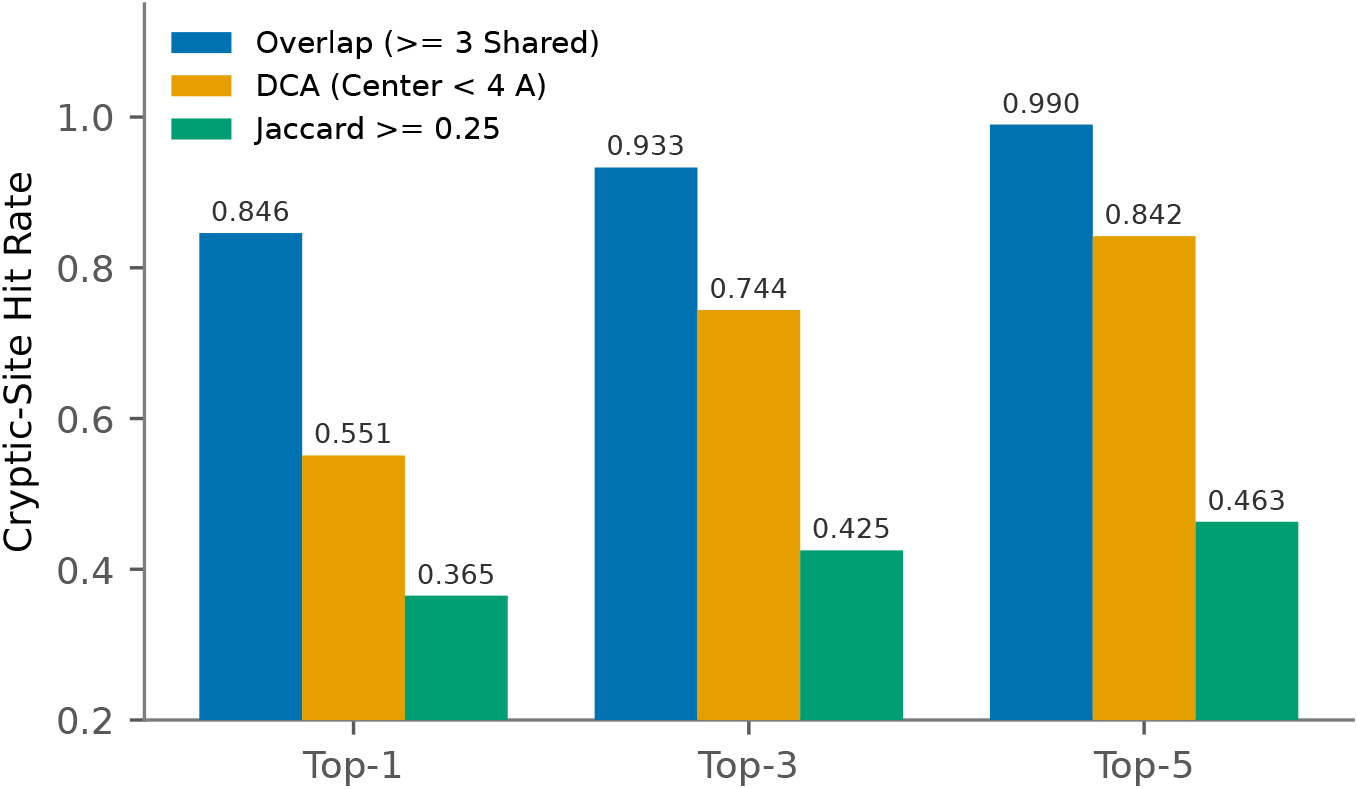
CryptoBank detection across rules. Cryptic-site hit rate on a 286-chain subset of CryptoBank at ranks 1, 3 and 5 under the overlap, DCA and Jaccard rules.

### 5.2 Per-Residue Detection and the Exact Residue Set

Per-residue detection reaches an AUC of 0.8465. Defining the pocket as a set is more challenging than simply locating it, as reflected in the results: under the stricter Jaccard rule, the detector recovers the site at top-1 0.597 and top-5 0.628. Figure 2 presents the overlap, DCA, and Jaccard rules side by side; the difference between the more lenient overlap rule and the stricter Jaccard rule primarily reflects how well the predicted pocket matches the precise shape of the annotated site, rather than whether the correct region was identified. This distinction is why the rules are reported separately and not interchanged.

## 6. Held-Out Generalization

### 6.1 WRN Helicase, Held Out of Training

WRN helicase presents a challenging and well-studied test case because its allosteric site is cryptic and is defined only by inhibitor-bound structures. To ensure a rigorous evaluation, we removed the WRN chain (8PFP) from training. After this removal, the closest remaining training chain shares only 22.5% sequence identity, which is below the commonly accepted threshold for reliable sequence-based inference and can be verified through a leakage audit using public sequence records. The allosteric site is then recovered at rank 1 on all three independent apo crystal forms (6YHR, 9MJT, 9S1A). On the 6YHR form the rank-1 pocket covers 18 of 22 annotated residues, including C727 (Figure 4); residue-level recovery is lower on the more incompletely resolved 9S1A form, so the result we rely on is the pocket-level rank-1 recovery on all three forms rather than a single residue-overlap figure. The recovered pocket is the binding site of the clinical WRN inhibitor HRO761 (PDB 8PFO): in that complex the compound contacts C727 and six further residues of the site, none of which the detector sees, since it reads only the static apo backbone with every inhibitor-bound structure removed from training. The same pocket, and the same 6YHR apo form, feature in recent structure-based modeling of WRN by the Boltz team [14]; that work reads the site from the bound complex (8PFO), whereas here it is recovered from the apo backbone alone, with 8PFO, 8PFP, and 7GQU all held out of training. Since WRN’s ligandable surface is essentially one clustered region, we report only the two sites with genuine structural support, the allosteric site (28 residues across 19 structures) and the nucleotide site (12 residues across 9 structures), and do not count the region’s several single-structure fragment hits as independent discoveries.

**Figure 4:**
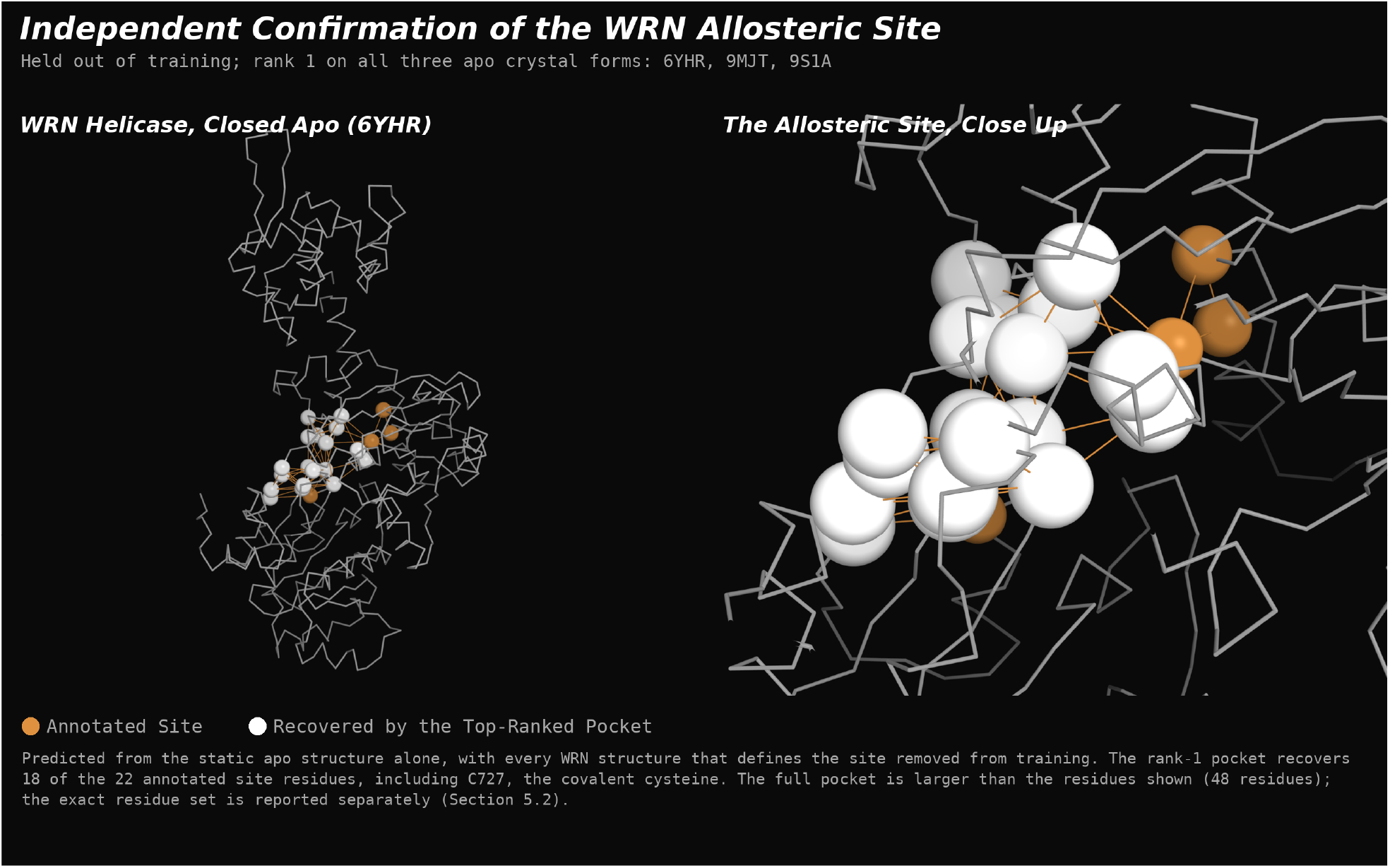
WRN allosteric site, held out. WRN helicase closed apo form 6YHR: the annotated allosteric site, with the residues recovered by the rank-1 pocket in white and those not recovered in orange. Predicted from the static apo structure alone, with every WRN structure that defines the site removed from training; the rank-1 pocket recovers 18 of 22 site residues, including C727, the covalent cysteine. The full pocket is larger than the recovered residues shown; the exact residue set is reported separately (Section 5.2).

### 6.2 Any Structure, Including mmCIF-Only

Because the detector requires only apo coordinates, it can operate on structures that are not present in any sequence database, which is essential for analyzing genuinely novel targets. Additionally, it is compatible with the current data format: since all depositions from 2025 onward are distributed exclusively as mmCIF files, the pipeline is designed to read mmCIF directly. As a result, recent targets are processed correctly, whereas tools limited to PDB format would fail to handle these entries.

## 7. Accuracy Scales with Data

The accuracy of the detector rises with the size of the training corpus, and the curve has not flattened even at the largest corpus we tested (Figure 5). Mean pocket Jaccard rises each time the number of training chains doubles, so the route to a better detector is more measured cryptic examples rather than a change of readout. The results above likely represent a lower bound that the same method should surpass as the corpus grows.

**Figure 5:**
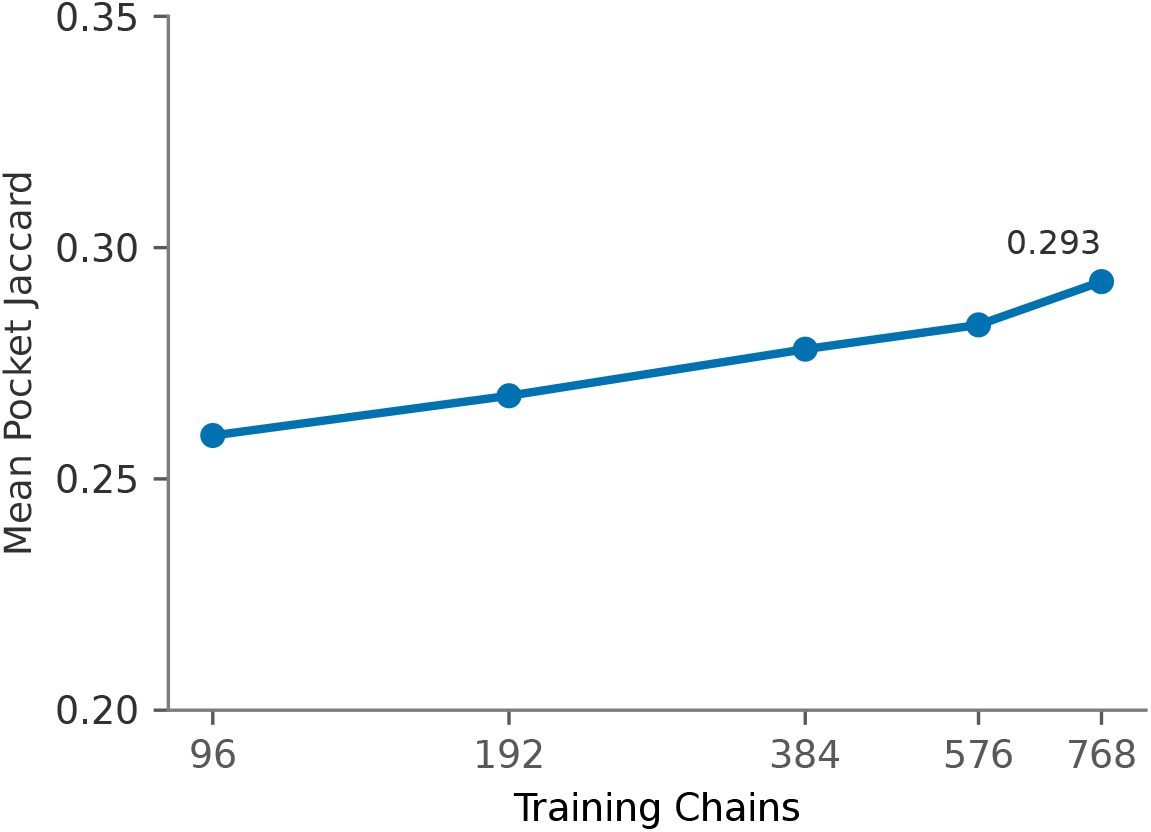
Accuracy scales with data. Mean pocket Jaccard against training-set size on a log-2 axis; the curve is still rising at the largest corpus.

## 8. Comparison with Baselines

### 8.1 Detection Baselines

Lacuna [7] reports two settings on the CryptoBench test fold, both at top-5 and both under its own evaluation rule (Jaccard at least 0.25 or a 4 angstrom centroid match): 0.556 with its default learned ranker, and 0.661 once an optional protein-language-model-assisted ranker is included. Under that same rule, our detector reaches 0.697 at top-1, surpassing Lacuna’s top-5 settings, and 0.840 at top-5; our numbers are measured on the full CryptoBench test split while Lacuna reports on a 180-structure subset of it, so the populations are close but not identical. Our rank-1 pocket alone exceeds what Lacuna reaches with five ranked candidates, its language-model ranker included.

Additionally, the two methods differ in their inputs: Lacuna generates and scores a conformational ensemble for each target, with its higher performance relying on a language-model ranker, whereas our detector analyzes a single static apo structure without conformational sampling or a protein language model at inference. Achieving ensemble-plus-language-model performance from a single static structure underscores the capability of the world model, which encodes where a cryptic pocket will open within the latent representation of the closed state, eliminating the need for external generation or language models. As in all comparisons, each method is evaluated under its own rule and denominator, ensuring that stricter and looser criteria are not inappropriately compared.

This comparison also highlights the core challenge of the task. Lacuna’s own analysis distinguishes between two aspects: coverage, whether the cryptic site appears among the top candidates, and conversion, whether the true site is ranked first. While ensemble detection methods largely address coverage, they often leave conversion unresolved [11]. This distinction is evident in our results as well: a top-5 overlap rate of 0.952 against a top-1 of 0.848 on CryptoBench (Section 5.1), and the world-model latent contributes most at top-1. Because the same per-residue score that surfaces the region also ranks the candidates, the true site is placed first, rather than simply being present among the top candidates. This ability to rank, which is a known limitation for ensemble detectors, is directly addressed by reading the site from a learned state instead of scoring geometry across sampled conformers.

Against single-structure detectors evaluated on the same CryptoBench test set, the margin is wider still. At the DCA center-distance rule and top-1, our detector reaches 0.649, against 0.417 for P2Rank [4], 0.323 for DeepPocket [13], 0.294 for PocketMiner [12], and 0.258 for fpocket [3] (Figure 6). Each of these reads one structure, as ours does, but scores geometry or learned features on the closed state rather than a world-model state, and that difference is reflected in the gap in the figure.

**Figure 6:**
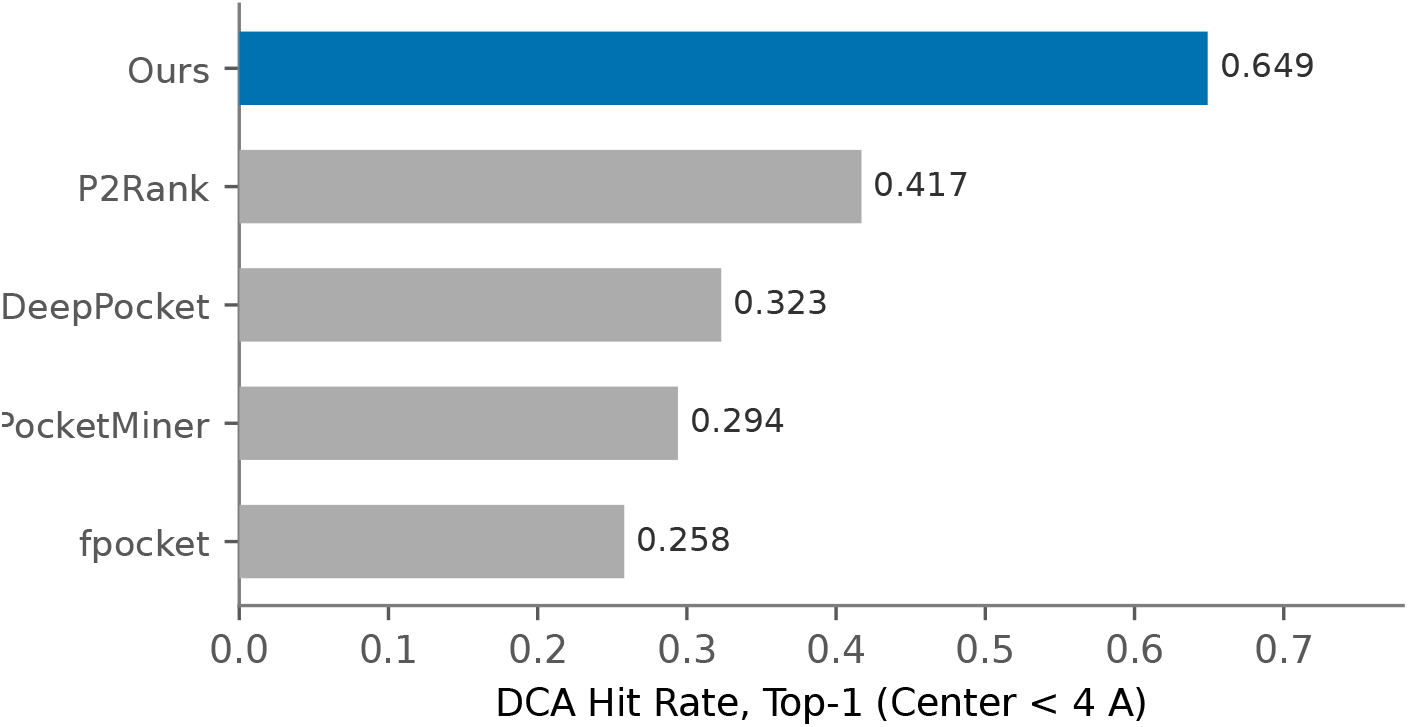
Single-structure baselines. Cryptic-site hit rate at the DCA center-distance rule and top-1 on CryptoBench, for our detector and four single-structure methods. All read one static structure; none samples conformations.

### 8.2 Direct Detection Versus Co-Folding

Co-folding engines such as OpenDDE [8] do not detect a pocket directly; they reach it only by predicting a bound complex, which means first finding a ligand that opens the site and reading the pocket off that prediction. Our detector predicts the pocket directly from the apo coordinates, with no ligand or bound complex required, so recovering a cryptic pocket is a first-class output of the method rather than a byproduct of solving the harder ligand-binding problem.

We make this concrete on 190 held-out cryptic targets absent from both training pools, scoring OpenDDE’s predicted contact residues and our own pockets under the same pocket-level rule (Figure 7). At our top-5, no target OpenDDE finds is one we miss: every pocket co-folding recovers, direct detection recovers too, plus 135 more, for a union of 181 of 190, so co-folding’s pocket-level outcomes are a subset of ours. At our top-1, we find 119 that OpenDDE does not, another 44 that OpenDDE also finds, and OpenDDE alone finds 2; 25 are found by neither method. We keep those two reverse cases visible rather than round them away. Direct detection is thus the general capability, and co-folding recovers a ligand-dependent slice of it.

**Figure 7:**
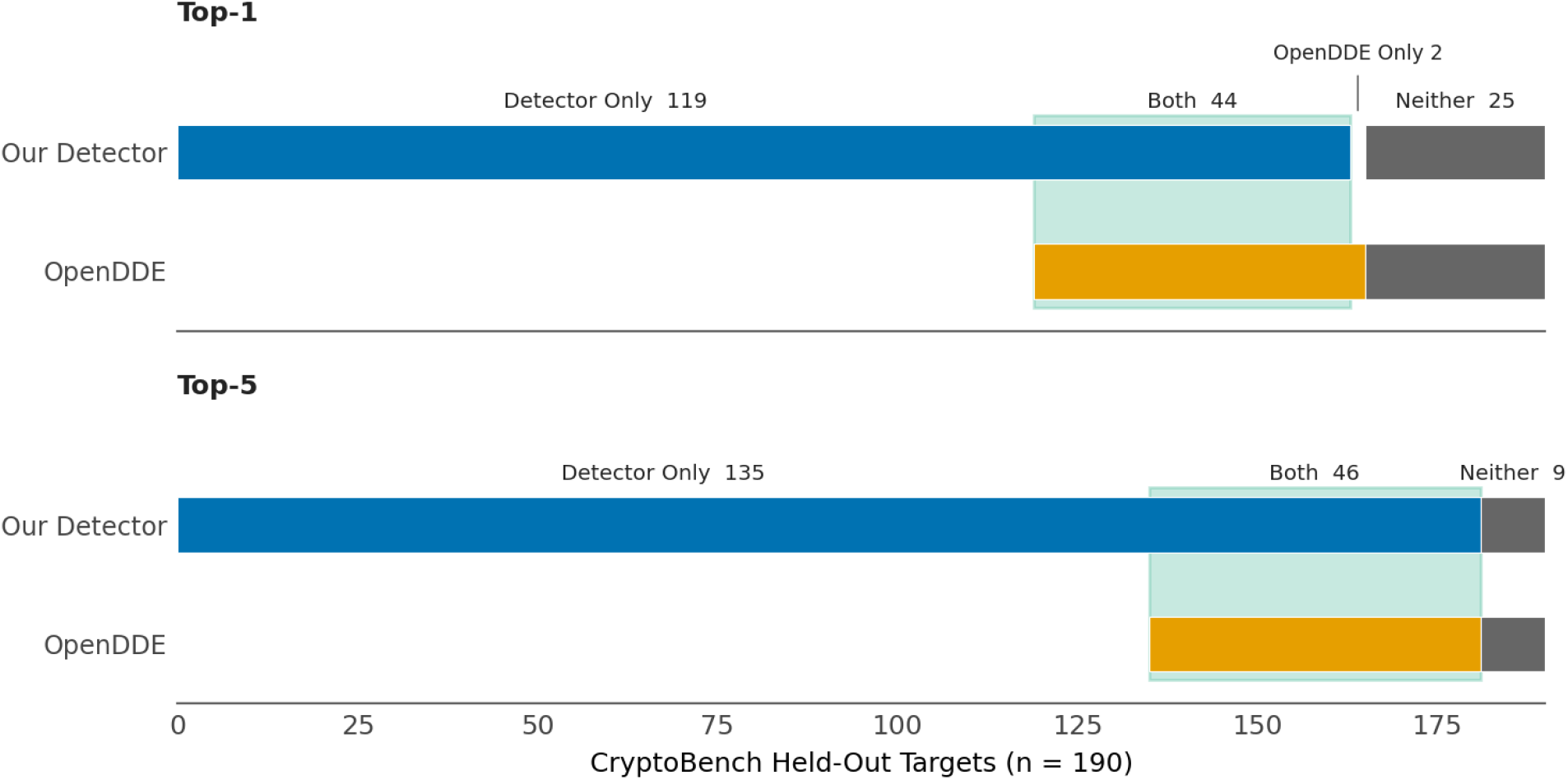
Direct detection versus co-folding. The 190 held-out cryptic targets decomposed by which method finds each, at our top-1 and top-5, scored identically at the pocket level. At top-5 no target OpenDDE finds is missed by us, so co-folding’s recovered pockets are a subset of ours.

## 9. Discussion and Scope

The results presented above are subject to several important limitations, each of which also highlights areas where the method could be improved.

A held-out chain does not equate to holding out an entire protein. A model that performs well on corpus-test data may not necessarily generalize to truly novel proteins, and this distinction is critical for prospective users. For this reason, we perform holdouts at the protein level throughout and use WRN, a target with no close homologs remaining in the training set, as a more stringent test case.

Identifying the location of a cryptic site and precisely mapping its residue set are distinct tasks, and the detector performs better at the former: overlap-rule metrics are high, whereas strict Jaccard metrics are lower. For the intended application, nominating the location of a cryptic pocket on otherwise intractable targets, localization is the most relevant measure, and the residue-set gap is expected to improve as more data become available.

Detection of cryptic sites is distinct from improving downstream design performance. While the world model aids in identifying pockets, it does not inherently enhance subsequent docking or co-folding design steps, where the primary challenge often lies in the docking assay rather than pocket identification; we do not claim otherwise. The WRN case further illustrates this limitation: under the strict center-distance metric, WRN falls into the more challenging category, being recovered in terms of pocket rank rather than precise center location. Therefore, both metrics should be considered together when interpreting results.

Finally, co-folding reaches cryptic pockets only through the harder ligand-binding route, so its pocket-level outcomes are largely a subset of ours, obtained indirectly, and at top-5 our detector misses none of OpenDDE’s hits. The residual value of a bound-complex view is confined to the few top-1 targets OpenDDE uniquely resolves, while direct detection is the general capability and covers the targets where no complex can yet be predicted.

## 10. Conclusion

A structural world-model latent enables detection of cryptic pockets directly from the apo structure of any protein, without the need to generate the open state as required by ensemble and co-folding methods. This approach generalizes to targets held out from training. The information about the site that will open is already encoded in the closed-state latent representation, eliminating the need for external data or processing. Furthermore, as the size of the training corpus increases, the accuracy of the detector improves, making prospective cryptic-site nomination on otherwise intractable targets both feasible now and increasingly effective as more data become available.

## Availability

The public corpora used are described in Section 4, and the evaluation protocol in that section defines every reported number. The representation and its training procedure are proprietary and are not released.

## Broader Impacts

Cryptic-site detection widens the set of proteins that can be considered for small-molecule intervention, including targets that look undruggable from their resting structure, which is broadly beneficial for programs against currently intractable disease targets. The capability that nominates a site does not validate it: a predicted pocket is a hypothesis for experimental follow-up. It does not show that a ligand exists or that engaging the site is safe or selective, and nominations should still pass the usual target-validation and medicinal-chemistry steps before any commitment. And because the detector reads only structure, it inherits the coverage and biases of the available structural data, so predictions on structurally under-represented families warrant correspondingly more caution.

